# Deep learning reveals many more inter-protein residue-residue contacts than direct coupling analysis

**DOI:** 10.1101/240754

**Authors:** Tian-ming Zhou, Sheng Wang, Jinbo Xu

**Author notes:** co-first authors (Tian-ming Zhou, Sheng Wang).

## Abstract

Intra-protein residue-level contact prediction has drawn a lot of attentions in recent years and made very good progress, but much fewer methods are dedicated to inter-protein contact prediction, which are important for understanding how proteins interact at structure and residue level. Direct coupling analysis (DCA) is popular for intra-protein contact prediction, but extending it to inter-protein contact prediction is challenging since it requires too many interlogs (i.e., interacting homologs) to be effective, which cannot be easily fulfilled especially for a putative interacting protein pair in eukaryotes. We show that deep learning, even trained by only intra-protein contact maps, works much better than DCA for inter-protein contact prediction. We also show that a phylogeny-based method can generate a better multiple sequence alignment for eukaryotes than existing genome-based methods and thus, lead to better inter-protein contact prediction. Our method shall be useful for protein docking, protein interaction prediction and protein interaction network construction.

## Introduction

Very few proteins perform their functions alone. Instead, most proteins function by interacting with others to form complexes and/or protein-protein interaction (PPI) networks[1–15]. To understand PPIs at structure level, their 3D structures may be needed. Although significantly improved, solving the 3D structure of a PPI or protein complex by experimental techniques is still very challenging, even if its constituent proteins have known 3D structures[16, 17]. There is still little 3D structure information for a big percentage of currently known PPIs in bacteria, yeast and human[18]. Therefore, computational methods are needed to elucidate the 3D structure of a PPI or complex from its sequence.

Predicting the 3D structure of a PPI or complex is very challenging. Here we focus on an important intermediate step that predicts inter-protein residue-residue contacts between a pair of putative interacting proteins (i.e., inter-protein contact prediction). Intra-protein contact prediction has demonstrated its effectiveness on folding a single protein chain [19–24]. It is also reported that inter-protein contact prediction is useful for the 3D structure modeling of a PPI or protein docking[16, 25, 26]. A few computational methods have been developed to predict protein-level interaction, including phylogenetic profiling[27], force field methods[28], genomic co-localization [29, 30] and others[31–33]. Nevertheless, these methods can only predict if two proteins interact or not, but not which residue pairs may interact or form a contact. Here our focus is to predict residue-level interactions (i.e., contacts) across two proteins.

Co-evolution analysis or more specifically direct-coupling analysis (DCA) is popular for intra-protein contact prediction[19, 34–39] and has also been applied to inter-protein contact prediction [16, 25, 37, 40, 41]. DCA works by identifying co-evolved residues from a multiple sequence alignments (MSA) of sequences homologous to a protein under prediction. Although popular, DCA has low prediction accuracy when we cannot find a large number of sequence homologs for a protein under prediction. This problem becomes more serious for inter-protein contact prediction since it is challenging to find so many interlogs (i.e., interacting homologs) for an interacting protein pair. Because of this, currently DCA for inter-protein contact prediction mainly focuses on prokaryotes and mitochondria[16, 25] since it is relatively easy to find interlogs in prokaryotes, but does not fare well on eukaryotes with abundant paralogs because it is challenging to identify correct interlogs [42].

We have developed a deep learning (DL) method for intra-protein contact prediction[20, 21], which greatly outperformed DCA and obtained the highest F1 score in CASP12[22]. Our DL method not only works for soluble proteins, but also for membrane proteins even if trained by only non-membrane proteins[21]. Compared to pure DCA, our DL method needs much fewer sequence homologs to be effective because it makes use of contact occurrence patterns, in addition to co-evolution information, for contact prediction. Inspired by the success of our DL method and the fact that it does not need so many sequence homologs, this paper studies how well DL can work on inter-protein contact prediction, especially for eukaryotic species. To avoid overfitting, we do not train our DL model using any protein complex data (i.e., inter-protein contact maps). Instead, we still use our previous DL model trained by only individual protein chains (i.e., intra-protein contact maps) to predict inter-protein contacts.

To further improve accuracy, we propose a new phylogeny-based method to identify interlogs for a putative interacting protein pair, especially for eukaryotes in which some interacting genes may be far away from each other along genome. When coupled with DL, this new method works better on eukaryotes than genome distance-based methods employed by Baker[16] and Marks[25] (which are mainly developed for prokaryotes). Experimental results show that combining these two methods can even further improve prediction accuracy.

We have tested our DL method using two datasets adopted by other groups and a large number of interacting protein pairs extracted from 3DComplex[43]. Experimental results show that our DL method indeed outperforms pure DCA for inter-protein contact prediction. Our DL method not only works on prokaryotes, but also on eukaryotes and the other species. We have also observed that inter-protein contact prediction accuracy is not only highly related to the number of effective interlogs available for a protein pair under prediction, but also to the density of inter-protein contacts defined as the ratio between the number of inter-protein contacts and the total sequence length.

## Method

Following the definition in[16, 25], we say that two residues of a protein pair form an inter-protein contact if 1) they belong to two different protein chains; and 2) in the native 3D structure of a complex containing this protein pair, the minimum Euclidean distance between their heavy atoms is less than 6Å. We measure the density of inter-protein contacts across two proteins by *#contacts/L* where *#contacts* is the number of true contacts across two proteins and *L* is their total sequence length.

**Overview**. As shown in Fig. 1, given a pair of putative interacting proteins A and B for which we would like to predict inter-protein contacts, we first run HHblits[44] to find sequence homologs and build multiple sequence alignments (MSAs) for A and B, respectively. Then we employ two different strategies to concatenate MSA_A and MSA_B into a paired MSA consisting of only interlogs (i.e., interacting homologs). Let MSA_g and MSA_p be the paired MSAs resulting from genome-based and phylogeny-based methods (described below), respectively, from which we may use our DL method to predict two inter-protein contact maps, and then calculate their average as the final prediction.

**Figure 1.**
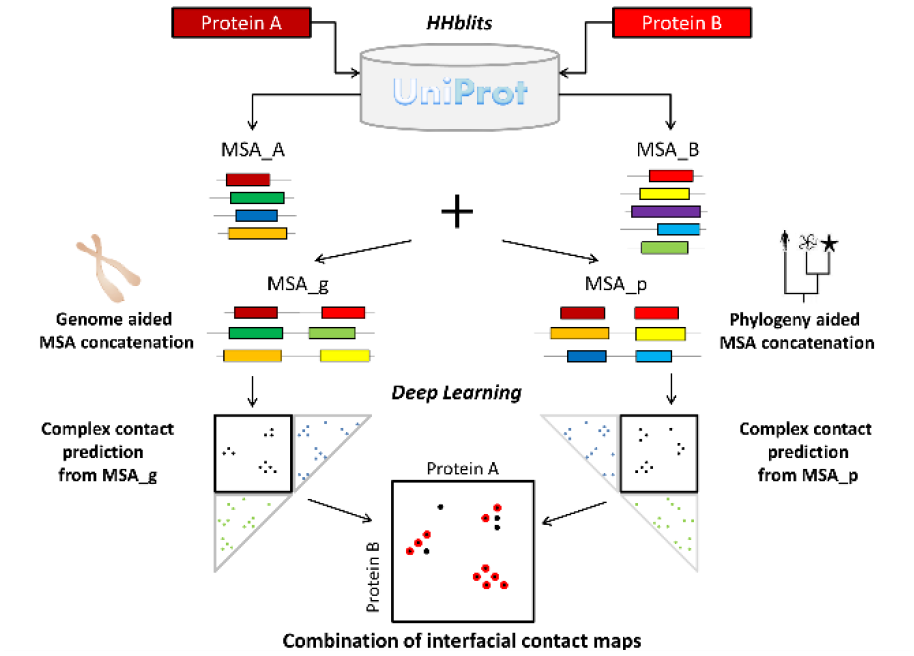
Schematic flowchart of our method.

### Building multiple sequence alignment (MSA) for a protein pair

We generate an MSA for an individual protein by running HHblits with 8 iterations and E-value=1E-20 to search through the UniProt20 HHM library dated in February 2016. The other parameters for HHblits are -maxfilt 100000000 -diff inf -all -neffmax 20. Let MSA_A and MSA_B denote the resultant individual MSAs for proteins A and B, respectively. We employ two strategies to concatenate MSA_A and MSA_B into a paired MSA, which consists putative interacting pairs homologous to the query pair. One is based upon genome information and the other utilizes phylogeny information. Not every protein in MSA_A and MSA_B can be paired. After concatenation, unpaired proteins are removed.

#### Concatenating MSAs by genomic distance

In prokaryotes and some eukaryotes, interacting genes are often co-located on the chromosome into operons[45], so we may tell if two proteins forming an interacting pair (i.e., interlog) or not by checking if their intergenic distance is less than a given threshold (we use 20 here). Both EVcomplex[25] and Baker[16] employed this strategy, but differed in some details. We may use a bipartite graph to describe the pairing relationship between proteins. That is, we treat each individual protein as a node and add one edge between two nodes (of two different MSAs) if their intergenic distance is less than 20. We may assign the inverse of the intergenic distance as weight to an edge. Then, we may pair two MSAs by finding the maximum matching in the graph[46]. Before doing so, we may also prune edges of which at least one endpoint does not have the minimum distance to the other. After pairing proteins in two MSAs, we remove the unpaired proteins and generate a paired MSA consisting of only putative interlogs, each being supposed to be homologous to the query protein pair.

Fig. 2 shows an example of this method. Here the left (red) and right (blue) MSAs contain 5 and 6 protein chains, respectively, and the intergenic distance threshold is set to 3. In total there are 6 contigs containing 6 copies of proteins (red blocks) in the left MSA and 8 copies of proteins (blue blocks) in the right MSA. Grey blocks are coding DNA sequences not belonging to these two MSAs. Based upon intergenic distance, 4 protein pairs can be formed, as shown by the black double arrows. For example, in contig 2, the intergenic distance between proteins 2 and B is 1, smaller than that between proteins 2 and A, so we pair proteins 2 and B together. In contig 4, the intergenic distance between proteins 4 and D is 4, larger than the threshold, so we do not pair them together.

**Figure 2.**
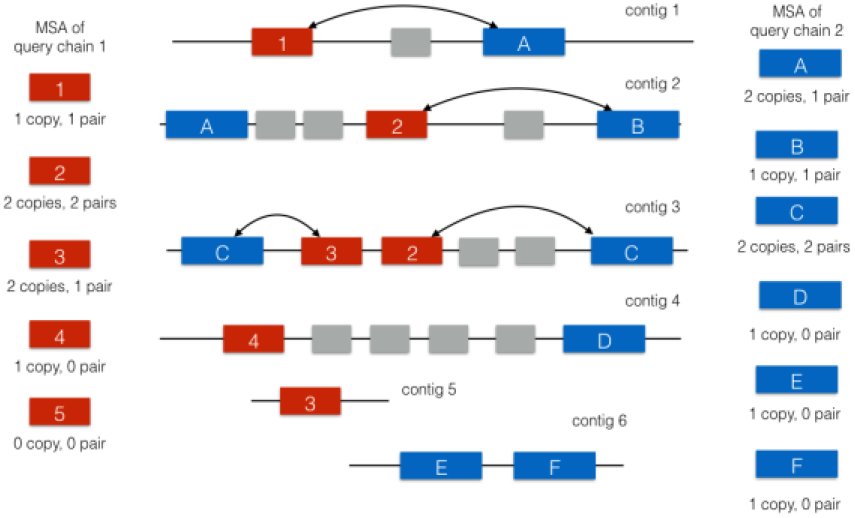
An example for genome-based MSA concatenation.

To calculate intergenic distance, we need map proteins (more specifically, coding DNA sequence (CDS)) to chromosome. To do so, we extract the UniProt IDs of a protein from the database and fetch the contig IDs from European Nucleotide Archive (ENA) via its XREF service[47]. Then we retrieve the contigs from ENA by their IDs and extract CDS lines that map UniProt IDs to their positions in contigs. Two CDSs are considered belonging to one gene if their spanning regions have an overlap of length at least one CDS.

#### Concatenating MSAs by phylogeny information

It is much more challenging to concatenate two individual MSAs for a protein pair in eukaryotes. This is because an individual MSA may contain abundant paralogs and two genes may interact with each other even if they are not close by genomic distance [40, 42]. Here we propose to concatenate two individual MSAs using phylogeny and homology information. First, we group proteins in each MSA by their species (or sub-species if possible) according to the phylogeny tree stored in the Taxonomy Database[48]. Then we sort proteins of a specific species/subspecies in each MSA by their sequence similarity (from high to low) to their respective query proteins. Let p_1_, p_2_, …, p_m_ be the sorted proteins of a specific species in one MSA and q_1_, q_2_, …, q_n_ be the sorted proteins of the same species in the other MSA. Then we pair p_i_ and q_i_ together where *i* ranges from 1 to the minimum of *m* and *n*.

Fig. 3 shows an example of this method. Each individual MSA contains 8 proteins (including the query). *Manis pentadactyla* and *Alligator sinensis* have no subspecies. *Ailuropoda melanoleuca* has two subspecies: *A.m.melanoleuca* and *A.m.qinlingensis*. Therefore, we may divide the proteins into 4 groups. *Manis pentadactyla* has only one protein in each MSA, so we simply pair its two proteins to form an interlog. *Alligator sinensis* has 2 and 3 proteins for queries 1 and 2, respectively. We sort them by sequence similarity and remove the protein which is the least similar to query 2 and pair the remaining proteins to form 2 interlogs. Similarly, we obtain 1 interlog for *A.m.qinlingensis* and 2 for *A.m.melanoleuca*. Finally, we obtain a paired MSA of 7 interlogs (including the query itself), as shown by the black double arrows.

**Figure 3.**
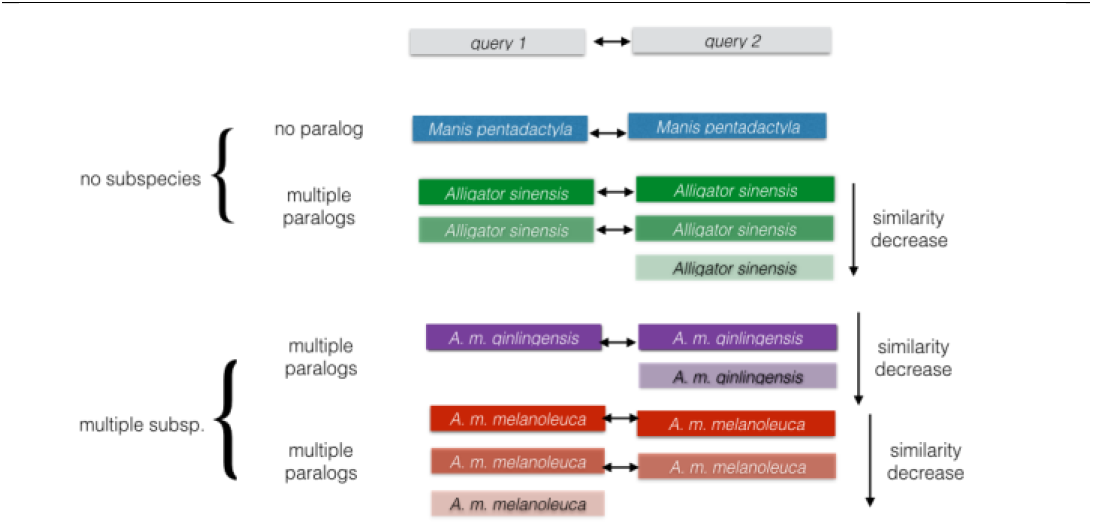
An example for phylogeny-based MSA concatenation.

#### Calculating the number of effective interlogs

Meff measures the amount of homologous information in a multiple sequence alignment (MSA). It can be interpreted as the number of non-redundant (or effective) sequence homologs in an MSA. Here we use 70% sequence identity as cutoff to determine if two interlogs are similar or not. Two interlogs are similar if and only if two proteins in one interlog are similar to their corresponding proteins in the other interlog. Let Si denote the number of interlogs (including itself) similar to interlog i. Then we may calculate Meff of an MSA as the sum of 1/Si over all the interlogs in this MSA.

### Protein features for a protein pair

For each protein pair under prediction, we derive two kinds of protein features (sequential and pairwise features) and then feed them into our DL model for contact prediction. Sequential features include protein sequence profile (i.e., position-specific scoring matrix), 3-state secondary structure[49] and 3-state solvent accessibility[50] predicted by our in-house tool RaptorX-Property[51]. We generate sequential features separately for two proteins in a pair and then concatenate them to form paired sequential features. That is, we treat a protein pair as a single protein by concatenating their sequences. Sequence profile is represented as a 2D matrix with dimension L×20 where L is the total protein length. Predicted secondary structure is represented as a 2D matrix with dimension L×3 (each entry is a predicted probability), so is the predicted solvent accessibility. Combining them together, we have a 2D matrix with dimension L×26 as the input sequential feature.

To calculate pairwise features, we feed a paired MSA to CCMpred[34], which outputs direct co-evolution strength between any two residues (i.e., columns) in the MSA. We also calculate mutual information and pairwise potential between any two columns in a paired MSA. See our previous work[20] for details.

### Deep learning (DL) for inter-protein contact prediction

As shown in Fig. S1 in Appendix, our DL model is formed mainly by two deep residual neural networks (ResNet)[52]. One is used to handle sequential features and the other pairwise features. The first ResNet conducts 1-dimensional (1D) convolutional transformation of sequential features to capture long-range sequential context of each residue in the query proteins. Its output is converted to a 2-dimensional (2D) matrix by an operation similar to outer product and then fed into the 2nd ResNet together with the original pairwise features. The 2nd ResNet conducts 2D convolutional transformation of its input to capture long-range 2D context of a residue pair. Finally, the output of the 2nd ResNet is fed into logistic regression, which predicts the probability of any two residues forming a contact. Please see our previous work [20, 22] for a detailed description of this DL model.

#### Combining contacts predicted from MSAs constructed by two strategies

For a protein pair, we build two paired MSAs using both genome- and phylogeny-based methods, and from the two MSAs predict two inter-protein contact maps, respectively. Then we may calculate their arithmetic average as the final prediction for this protein pair.

### Training and test datasets

We employed three testsets. (1) Baker’s dataset: a set of 32 protein pairs used by Baker to test a direct coupling (DCA) method Gremlin[16], 28 of which are from prokaryotes. This set includes protein pairs extracted from bacterial 50S ribosomal unit, the TRAP complex, tripartite efflux system, pyruvate formate lyase-activating enzyme complex and D-methionine transport system. (2) EVcomplex’s dataset[25]: a set of 86 protein pairs, in which 82 are from prokaryotes. (3) 3DComplex version 2.0[43]: 3067 non-redundant (40% sequence identity cutoff) complexes with experimentally-solved structures in PDB. Here we only consider heteromers, so finally obtained 4954 chain pairs in contact that involve 5296 protein chains. Among them, 841 and 178 pairs belong to human and E. coli., respectively; 1863 and 728 pairs belong to the other eukaryotes and prokaryotes, respectively. The remaining 1344 pairs belong to neither prokaryotes nor eukaryote (e.g. archaea and virus). In this set, 475 of protein pairs are too long and CCMpred cannot run them on GPUs due to memory limit, so we exclude them from consideration.

Our DL model is trained and validated using only individual protein chains (i.e., intra-protein contacts) in PDB25. That is, we did not train our DL model using any protein pairs (i.e., inter-protein contacts). A benefit of training by only protein chains is that there is no redundancy between our training data and the test data. To further verify this, we concatenate two proteins in a test pair to a single sequence and BLAST it against our training proteins. It turns out that BLAST cannot find any good alignment between the concatenated sequences and our training proteins. Please see our previous work on training our DL model[20].

### Evaluation and metrics

We evaluate our DL method by comparing it with a few pure DCA methods such as CCMpred[34], Gremlin[16] and EVcomplex[25]. In terms of methodology and accuracy, CCMpred, EVcomplex and Gremlin are not very different given the same input MSA. However, CCMpred runs much faster, so we focus more on it. We calculate the accuracy of the top 50, 20, 10, 5 and top L/k (k=5, 10, 20) predicted contacts where L is the total length of the two protein chains. The prediction accuracy is defined as the percentage of correctly predicted contacts among the top predictions.

## Result

### Prediction accuracy on the Baker’s and EVcomplex’s datasets

As shown in Tables 1 and 2, tested on these two datasets, our DL method (denoted as GCNN) greatly outperforms all pure DCA methods such as EVComplex (EVfold), Gremlin and CCMpred regardless of how many top predicted contacts are evaluated and no matter how the MSA is constructed. The results indicate that the genome-based method for MSA construction outperforms our phylogeny-based method on these datasets. This is because that most test protein pairs belong to prokaryotes and co-regulated genes in prokaryotes are often co-located on the genome into operons. Merging the contacts predicted from MSAs constructed by both phylogeny-based and genome-based methods does not help, partially because that the contacts predicted from phylogeny-based MSAs have much lower accuracy. Tables 1 and 2 confirm that CCMpred performs comparably to or better than EVfold, EVComplex and Gremlin, so in the following sections we will mainly compare our method with CCMpred.

**Table 1.**
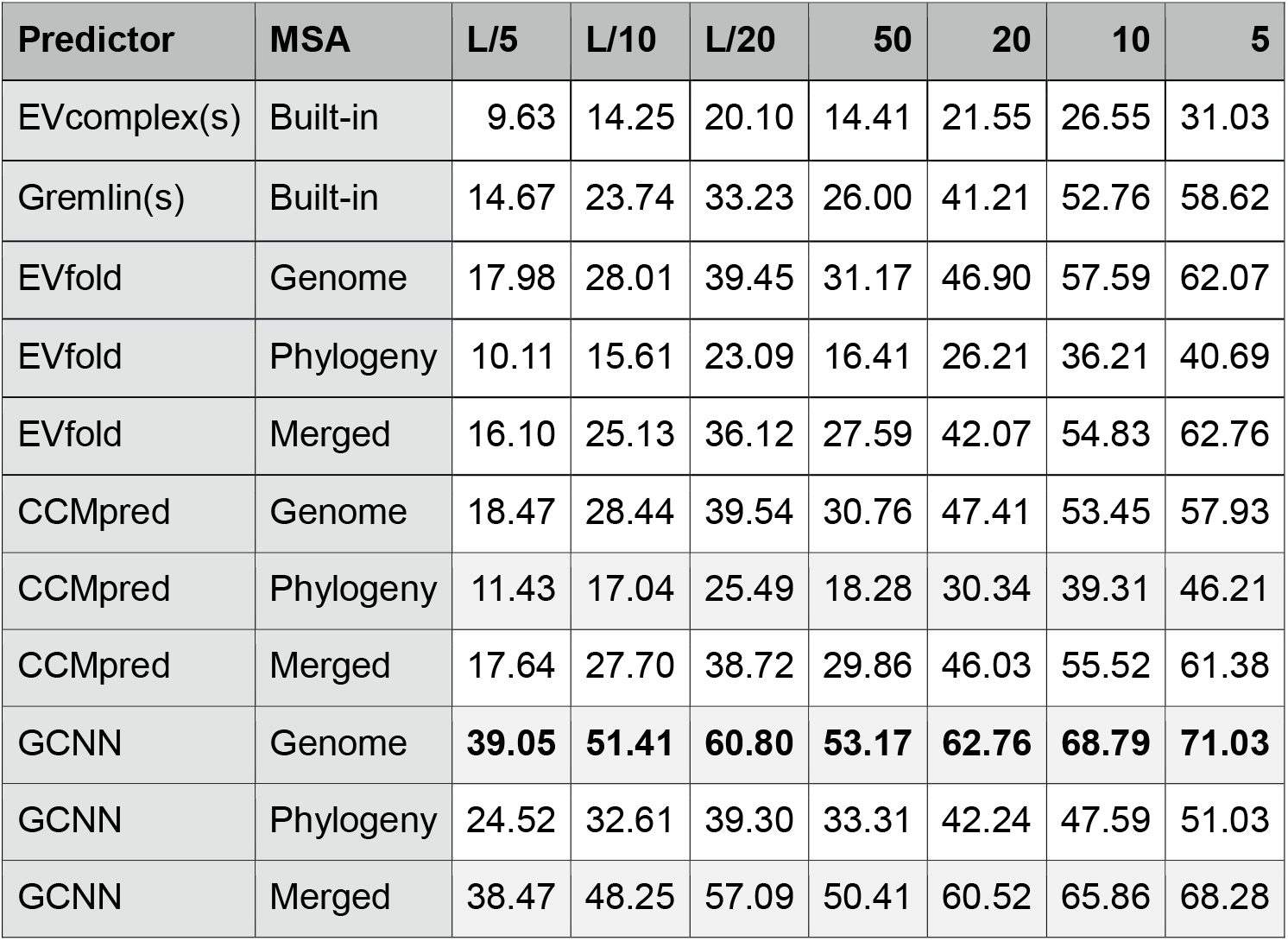
Inter-protein contact prediction accuracy (%) on Baker’s data. Our DL method is denoted as GCNN and (s) indicates a web server. EVfold is the same as EVcomplex, but run locally with our MSAs. “Genome” and “Phylogeny” denote genome- and phylogeny-based methods for MSA generation, respectively. “Merged” indicates prediction is merged from “Genome” and “Phylogeny”. Columns 3-9 show accuracy of top L/5, L/10, L/20, 50, 20, 10, and 5 predicted contacts.

| Predictor | MSA | L/5 | L/10 | L/20 | 50 | 20 | 10 | 5 |
| --- | --- | --- | --- | --- | --- | --- | --- | --- |
| EVcomplex(s) | Built-in | 9.63 | 14.25 | 20.10 | 14.41 | 21.55 | 26.55 | 31.03 |
| Gremlin(s) | Built-in | 14.67 | 23.74 | 33.23 | 26.00 | 41.21 | 52.76 | 58.62 |
| EVfold | Genome | 17.98 | 28.01 | 39.45 | 31.17 | 46.90 | 57.59 | 62.07 |
| EVfold | Phylogeny | 10.11 | 15.61 | 23.09 | 16.41 | 26.21 | 36.21 | 40.69 |
| EVfold | Merged | 16.10 | 25.13 | 36.12 | 27.59 | 42.07 | 54.83 | 62.76 |
| CCMpred | Genome | 18.47 | 28.44 | 39.54 | 30.76 | 47.41 | 53.45 | 57.93 |
| CCMpred | Phylogeny | 11.43 | 17.04 | 25.49 | 18.28 | 30.34 | 39.31 | 46.21 |
| CCMpred | Merged | 17.64 | 27.70 | 38.72 | 29.86 | 46.03 | 55.52 | 61.38 |
| GCNN | Genome | **39.05** | **51.41** | **60.80** | **53.17** | **62.76** | **68.79** | **71.03** |
| GCNN | Phylogeny | 24.52 | 32.61 | 39.30 | 33.31 | 42.24 | 47.59 | 51.03 |
| GCNN | Merged | 38.47 | 48.25 | 57.09 | 50.41 | 60.52 | 65.86 | 68.28 |

**Table 2.** Inter-protein contact prediction accuracy (%) on EVComplex’s data. See Table 1 for more explanation.

| Predictor | MSA | L/5 | L/10 | L/20 | 50 | 20 | 10 | 5 |
| --- | --- | --- | --- | --- | --- | --- | --- | --- |
| Gremlin(s) | Built-in | 5.22 | 8.16 | 11.51 | 7.16 | 12.03 | 16.05 | 20.47 |
| EVfold | Genome | 6.08 | 9.73 | 14.11 | 9.30 | 15.87 | 21.16 | 26.74 |
| EVfold | Phylogeny | 4.22 | 5.81 | 8.24 | 5.21 | 8.20 | 11.74 | 14.88 |
| EVfold | Merged | 5.58 | 8.56 | 12.71 | 8.05 | 13.20 | 18.49 | 24.19 |
| CCMpred | Genome | 6.05 | 9.01 | 13.81 | 8.35 | 13.95 | 19.77 | 23.72 |
| CCMpred | Phylogeny | 3.95 | 6.16 | 8.89 | 4.65 | 7.97 | 11.86 | 15.81 |
| CCMpred | Merged | 5.82 | 8.88 | 13.26 | 7.88 | 13.55 | 18.95 | 24.65 |
| GCNN | Genome | **13.70** | **18.18** | **22.58** | **17.51** | **22.79** | **26.51** | **30.00** |
| GCNN | Phylogeny | 8.81 | 12.51 | 16.40 | 10.70 | 15.99 | 19.19 | 22.09 |
| GCNN | Merged | 12.93 | 17.74 | 22.41 | 16.47 | 22.38 | 25.23 | 27.44 |

### Prediction accuracy on 3DComplex

Fig. 4 shows the top L/10 inter-protein contact prediction accuracy of the data in 3DComplex by our DL method and CCMpred on MSAs derived from two different strategies. One dot in the figure represents one test protein pair. Test protein pairs are colored by their species. Meanwhile, “N/A” indicates all species other than eukaryotes and prokaryotes. Fig. 4 shows that on most test protein pairs, our DL method outperforms CCMpred by a lot regardless of their species and how their MSAs are constructed.

**Figure 4.**
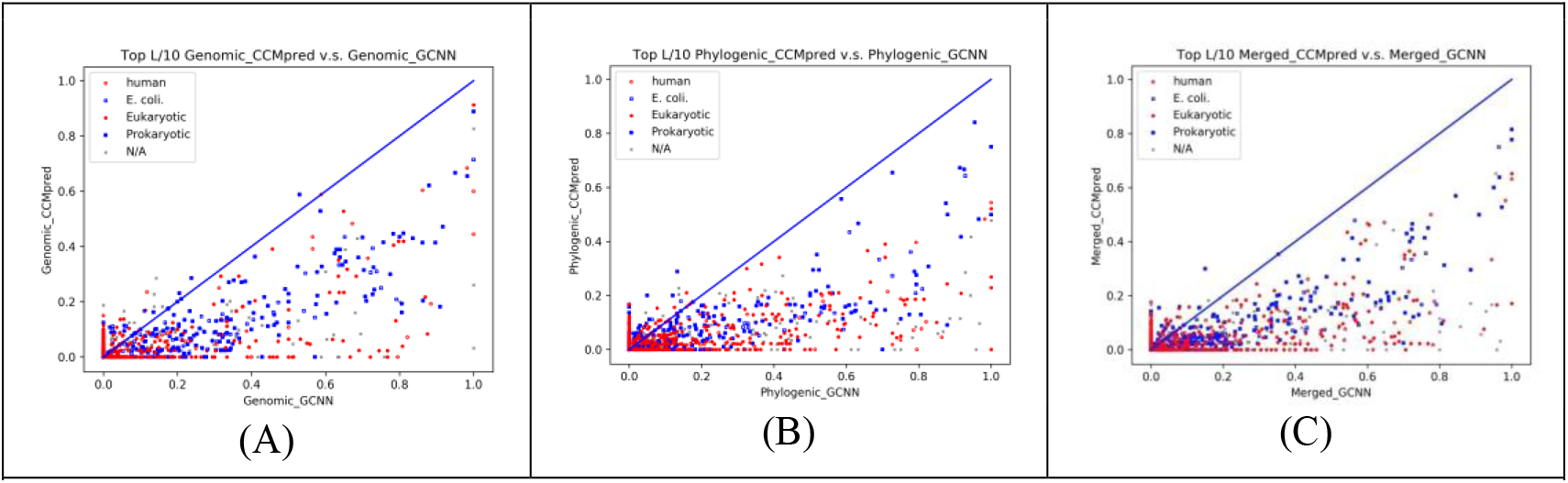
The top L/10 prediction accuracy by our DL method and CCMpred on the protein pairs extracted from 3DComplex. Each dot represents one test protein pair. (A) and (B) show the prediction accuracy of our DL method (X-axis) and CCMpred (Y-axis) on the MSAs derived by genome-based and phylogeny-based methods, respectively. (C) shows the accuracy of the contacts merged from the predictions represented by (A) and (B).

### Key factors impacting inter-protein contact prediction accuracy

We have observed that inter-protein contact prediction accuracy is related to two factors: the number of sequence homologs in an MSA (i.e., *Meff*) and inter-protein contact density measured by *#contacts/L* where *#contacts* is the number of native contacts across two proteins and *L* is their total length. We divide all the 3DComplex test protein pairs into groups by their *ln*(*Meff*) or contact density and then calculate the average accuracy in each group. Since for each protein pair we have two different MSAs, we use the geometric average of their *Meffs* to measure the *Meff* of one protein pair. Fig. 5 shows that the top *L/10* prediction accuracy increases with respect to both contact density and *Meff* and that our DL method greatly outperforms CCMpred regardless of *Meff* and contact density. We have also observed that when contact density is below 0.10, the prediction accuracy is low even if *Meff* is large. When contact density is larger than 0.25, the top L/10 accuracy increases faster when *Meff* is large than when *Meff* is small. This trend is similar for both our DL method and CCMpred.

**Figure 5.**
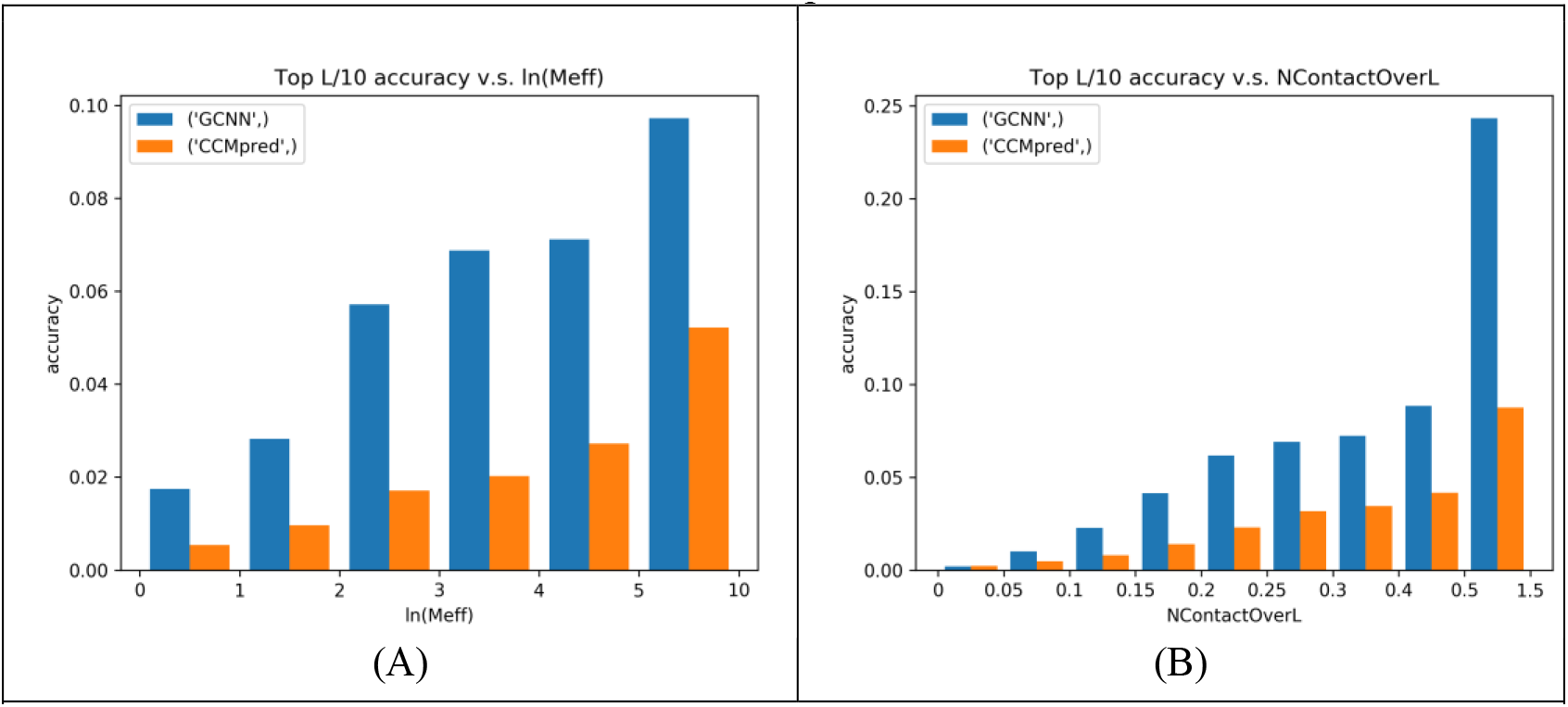
The average top L/10 accuracy (Y-axis) of our DL method and CCMpred with respect to (A) the number of sequence homologs measured by *Meff* and (B) contact density measured by #contacts/L. The test protein pairs are from 3DComplex. The last group in (A) has *ln(Meff)* between 5 and 10. The last group in (B) has contact density between 0.5 and 1.5.

Tables 3 and 4 further illustrate that prediction accuracy is related to contact density. Table 3 shows the top L/10 accuracy of our DL method and CCMpred on 125 randomly-selected protein pairs from 3DComplex, covering both prokaryotes and eukaryotes. Table 4 shows the top L/10 accuracy on 125 protein pairs from 3DComplex with the largest contact density. It shows that the prediction accuracy of our DL method and CCMpred is better by ~30% and ~10%, respectively, than those in Table 3, even when only top 5 or 10 predicted contacts are evaluated.

**Table 3.** Accuracy (%) on 125 randomly-selected protein pairs from 3DComplex. See Table 1 for more explanation.

| Predictor | MSA | L/5 | L/10 | L/20 | 50 | 20 | 10 | 5 |
| --- | --- | --- | --- | --- | --- | --- | --- | --- |
| CCMpred | Genome | 1.64 | 2.21 | 2.54 | 1.9 | 2.72 | 3.28 | 4.32 |
| CCMpred | Phylogeny | 1.71 | 1.82 | 2.31 | 1.71 | 2.32 | 2.88 | 3.84 |
| CCMpred | Merged | 1.91 | 2.3 | 2.94 | 2.06 | 3.00 | 4.08 | 5.12 |
| GCNN | Genome | 3.70 | 4.51 | 5.42 | 4.40 | 5.80 | 6.56 | 6.08 |
| GCNN | Phylogeny | 3.28 | 3.64 | 4.49 | 3.78 | 4.44 | 4.72 | 5.44 |
| GCNN | Merged | 4.32 | 5.09 | 5.73 | 4.96 | 6.28 | 7.04 | 7.68 |

**Table 4.** Accuracy (%) on 125 protein pairs from 3DComplex with the largest contact density. See Table 1 for more explanation.

| Predictor | MSA | L/5 | L/10 | L/20 | 50 | 20 | 10 | 5 |
| --- | --- | --- | --- | --- | --- | --- | --- | --- |
| CCMpred | Genome | 7.06 | 9.10 | 10.44 | 7.10 | 9.60 | 12.24 | 14.72 |
| CCMpred | Phylogeny | 7.58 | 9.20 | 12.37 | 8.26 | 11.00 | 15.20 | 18.08 |
| CCMpred | Merged | 9.16 | 12.08 | 15.31 | 10.02 | 13.95 | 18.20 | 19.60 |
| GCNN | Genome | 22.99 | 26.89 | 28.62 | 23.60 | 27.48 | 29.68 | 32.32 |
| GCNN | Phylogeny | 27.61 | 32.27 | 35.45 | 28.29 | 34.04 | 36.48 | 36.48 |
| GCNN | Merged | 30.02 | 34.38 | 36.83 | 30.69 | 35.44 | 37.76 | 38.08 |

### Phylogeny-vs. Genome-based methods for MSA construction

The phylogeny- and genome-based methods for MSA construction are complementary to each other. In general, for prokaryotic species, genome-based method works better and for eukaryotes, our phylogeny-based method works better. As shown in Tables 5 and 6, evaluated on 125 (50) test protein pairs (in 3DComplex) from all prokaryotic species (E. coli) with the largest contact density (i.e., *#contacts/L*), both our DL method GCNN and CCMpred have better prediction accuracy when the MSAs are derived from genome-based methods. Further, merging the predicted contacts from two types of MSAs yield higher accuracy on the 125 test protein pairs.

**Table 5.**
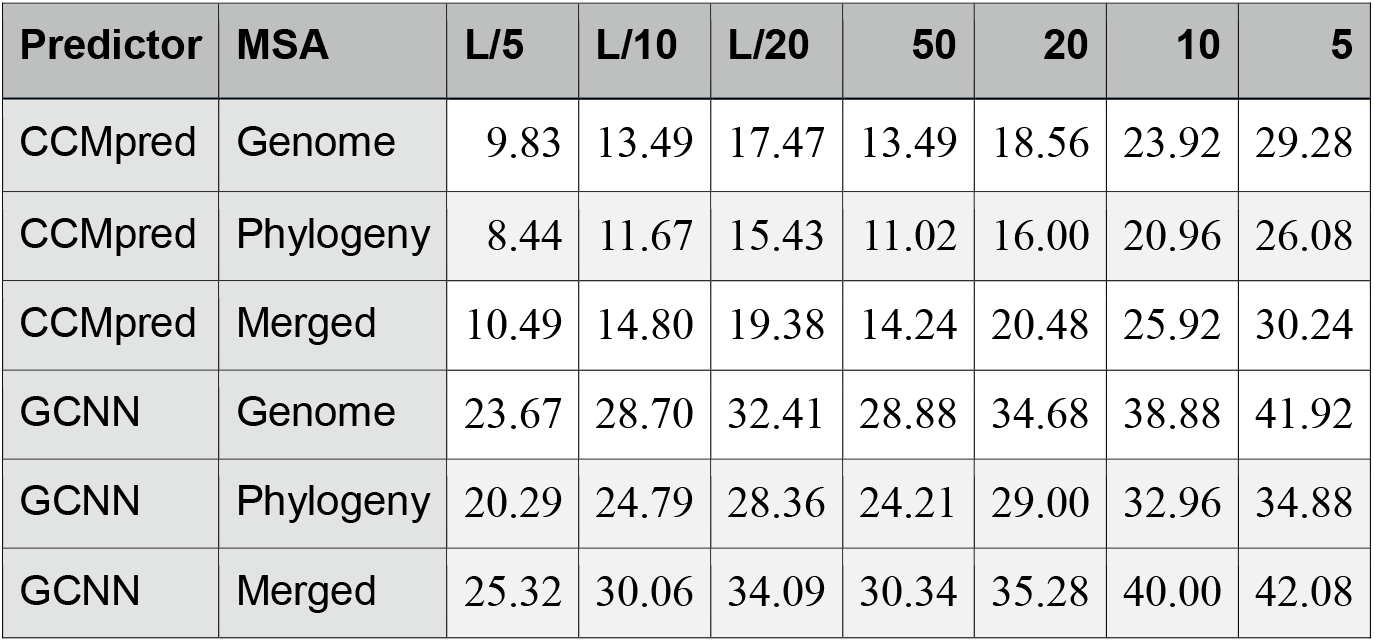
Inter-protein contact prediction accuracy (%) on 125 protein pairs from prokaryotic species in 3DComplex with the largest #contacts/L.

**Table 6.** Inter-protein contact prediction accuracy (%) on 50 protein pairs from E. coli in 3DComplex with the largest #contacts/L.

| Predictor | MSA | L/5 | L/10 | L/20 | 50 | 20 | 10 | 5 |
| --- | --- | --- | --- | --- | --- | --- | --- | --- |
| CCMpred | Genome | 8.73 | 12.24 | 17.20 | 11.68 | 17.30 | 22.40 | 31.20 |
| CCMpred | Phylogeny | 6.54 | 9.29 | 12.34 | 8.00 | 12.50 | 19.00 | 25.20 |
| CCMpred | Merged | 8.32 | 11.56 | 15.17 | 10.72 | 15.90 | 23.40 | 30.40 |
| GCNN | Genome | 22.98 | 27.59 | 31.85 | 27.64 | 35.70 | 39.60 | 39.20 |
| GCNN | Phylogeny | 18.70 | 23.99 | 29.82 | 22.84 | 30.50 | 34.40 | 37.20 |
| GCNN | Merged | 23.74 | 29.53 | 36.33 | 29.12 | 37.40 | 44.80 | 45.60 |

As shown in Tables 7 and 8, evaluated on 125(50) test protein pairs from eukaryotes (Human) with the largest contact density, the predicted contacts from the MSAs derived from our phylogeny-based method has better accuracy when coupled with our DL method. On both testsets, merging the predicted contacts from two types of MSAs yield higher accuracy than any single type of MSAs. The results in these 4 tables also confirm that our DL method greatly outperforms the pure co-evolution method such as CCMpred. Fig. S2 in Appendix also shows the contact prediction accuracy of individual protein pairs in 3DComplex resulting from these two MSA building methods.

**Table 7.** Inter-protein contact prediction accuracy (%) on top 125 chain pairs from eukaryotic species in 3DComplex with the largest #contacts/L ratio.

| Predictor | MSA | L/5 | L/10 | L/20 | 50 | 20 | 10 | 5 |
| --- | --- | --- | --- | --- | --- | --- | --- | --- |
| CCMpred | Genome | 4.58 | 6.28 | 7.95 | 4.50 | 5.88 | 7.52 | 8.80 |
| CCMpred | Phylogeny | 5.07 | 6.28 | 9.05 | 5.62 | 7.12 | 9.36 | 12.96 |
| CCMpred | Merged | 6.46 | 8.81 | 10.58 | 7.15 | 9.08 | 11.84 | 15.04 |
| GCNN | Genome | 15.17 | 17.85 | 18.77 | 14.42 | 17.92 | 19.20 | 20.96 |
| GCNN | Phylogeny | 21.78 | 24.99 | 25.93 | 22.75 | 26.68 | 26.64 | 26.24 |
| GCNN | Merged | 23.66 | 26.08 | 27.49 | 23.95 | 27.96 | 28.32 | 26.72 |

**Table 8.**
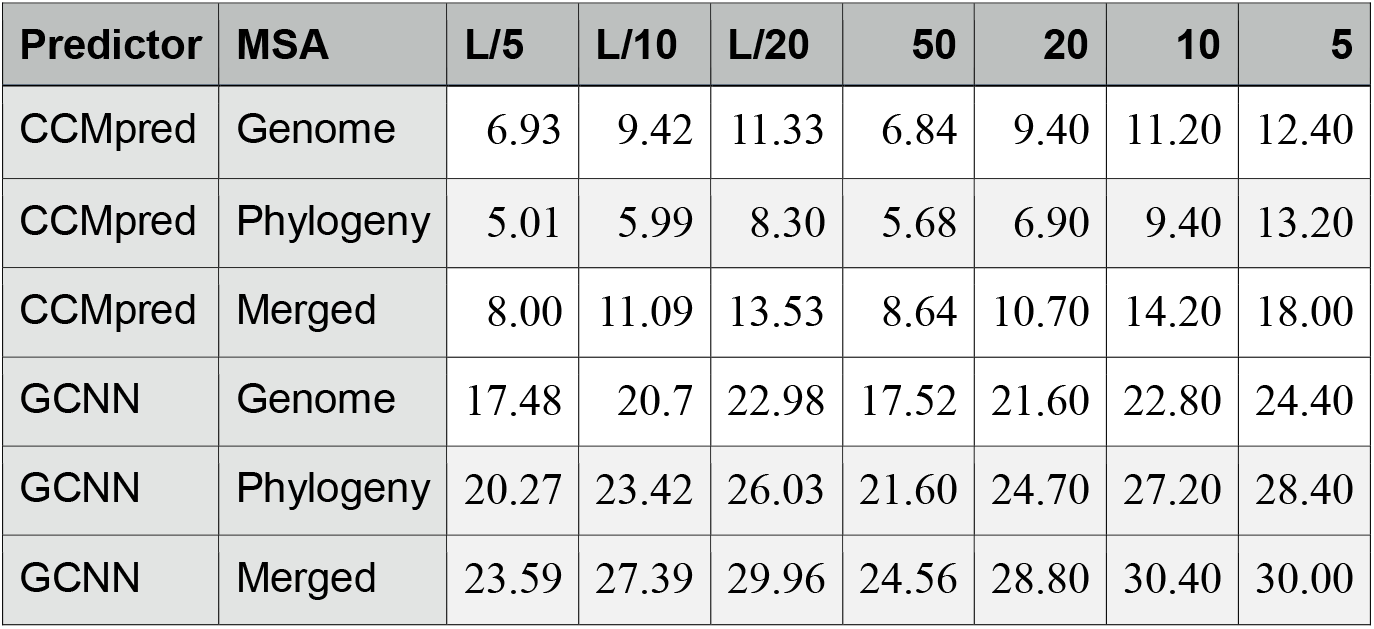
Inter-protein contact prediction accuracy (%) on 50 protein pairs from human species in 3DComplex with the largest #contacts/L ratio.

| Predictor | MSA | L/5 | L/10 | L/20 | 50 | 20 | 10 | 5 |
| --- | --- | --- | --- | --- | --- | --- | --- | --- |
| CCMpred | Genome | 6.93 | 9.42 | 11.33 | 6.84 | 9.40 | 11.20 | 12.40 |
| CCMpred | Phylogeny | 5.01 | 5.99 | 8.30 | 5.68 | 6.90 | 9.40 | 13.20 |
| CCMpred | Merged | 8.00 | 11.09 | 13.53 | 8.64 | 10.70 | 14.20 | 18.00 |
| GCNN | Genome | 17.48 | 20.7 | 22.98 | 17.52 | 21.60 | 22.80 | 24.40 |
| GCNN | Phylogeny | 20.27 | 23.42 | 26.03 | 21.60 | 24.70 | 27.20 | 28.40 |
| GCNN | Merged | 23.59 | 27.39 | 29.96 | 24.56 | 28.80 | 30.40 | 30.00 |

## Conclusion and Discussion

We have described the first deep learning (DL) method for inter-protein contact prediction that greatly outperforms existing pure co-evolution methods. Our approach works for both prokaryotes and eukaryotes. By contrast, existing co-evolution methods were mainly tested on prokaryotes. Our DL method is not trained by any existing protein complex data (i.e., inter-protein contact maps). Instead it is trained by only single-chain proteins (i.e., intra-protein contact maps), but still works for inter-protein contact prediction. The underlying reason is that both inter- and intra-protein contacts share similar physical-chemical properties[53] and contact occurrence patterns which are learned by our DL method. Nevertheless, the set of training intra-protein contact maps may not cover all possible contact occurrence patterns across two proteins. According to our experience in developing deep transfer learning for membrane protein contact prediction[21], we shall be able to further improve our DL method by training it using both intra- and inter-protein contact maps.

We have developed a phylogeny-based method for MSA construction, which outperforms genome-based methods on eukaryotes and provides information complementary to genome-based method. The combination of these two methods may lead to better prediction accuracy on eukaryotes. Although our DL method greatly outperforms pure co-evolution methods, the prediction accuracy of our method on eukaryotes is still not that high, maybe because the MSAs constructed by our phylogeny-based method is still suboptimal. We will continue to study better methods for MSA construction to further improve prediction accuracy.

Our experimental results show that prediction accuracy is related to inter-protein contact density, in addition to the number of interlogs (i.e., interacting homologs) available for a protein pair under prediction. Usually permanent PPIs has a larger inter-protein contact density than transient PPIs and thus, so our approach may have a better predictive effect for permanent PPIs and a better method is needed to predict contacts for transient PPIs.

Due to space limit, this paper focuses only on contact prediction, but not on building 3D structural models from predicted contacts. In an extended version we will describe how well our predicted contacts can be used to build 3D complex models of a PPI or to help improve protein docking. In this paper, we assume that we do not have the 3D structure models of the two constituent proteins of a putative interacting protein pair. Sometimes we may have native structures or good 3D models of individual proteins. In this case, we shall study how to further improve inter-protein contact prediction by making use of the known 3D structures of partner proteins.

## Acknowledgements

The authors are grateful to Ms. Anna Green of Harvard University for her help collecting results from the EVComplex web server.

## Appendix

## 1. Deep learning network architecture used for inter-protein contact prediction

**Figure S1.**
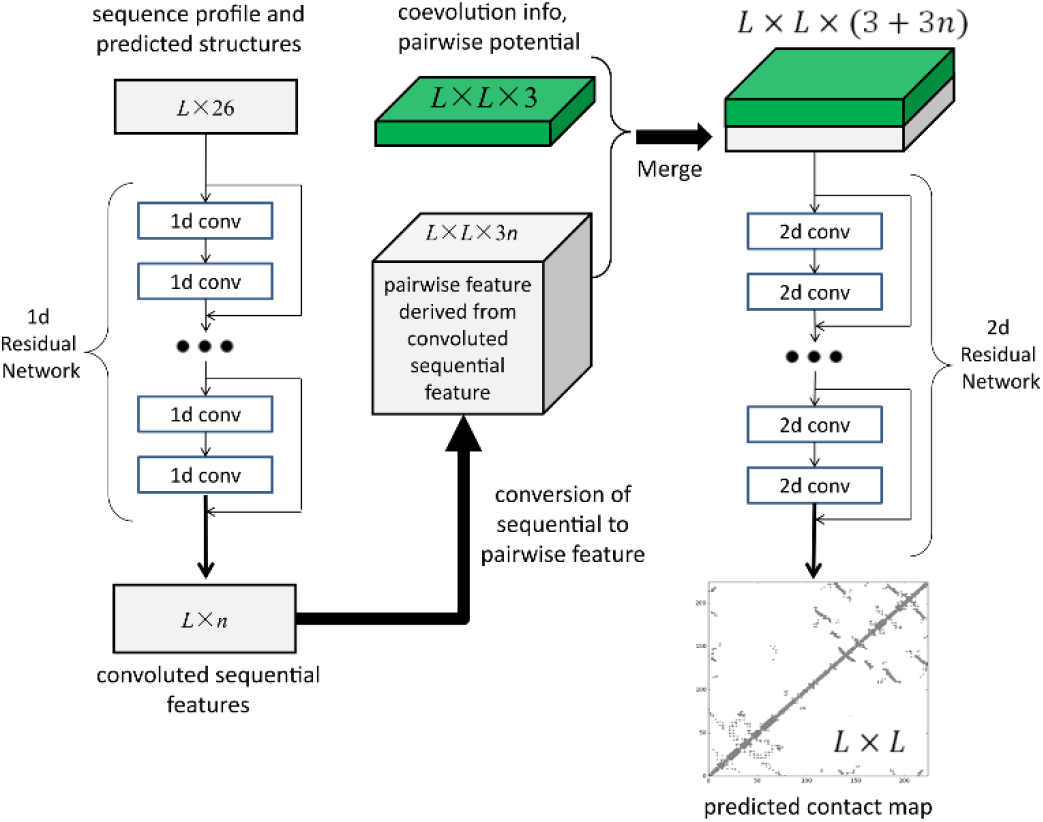
The network architecture used for contact prediction where L is the total protein length.

## 2. Comparison of phylogeny- and genome-based methods for MSA construction in terms of prediction accuracy

**Figure S2.**
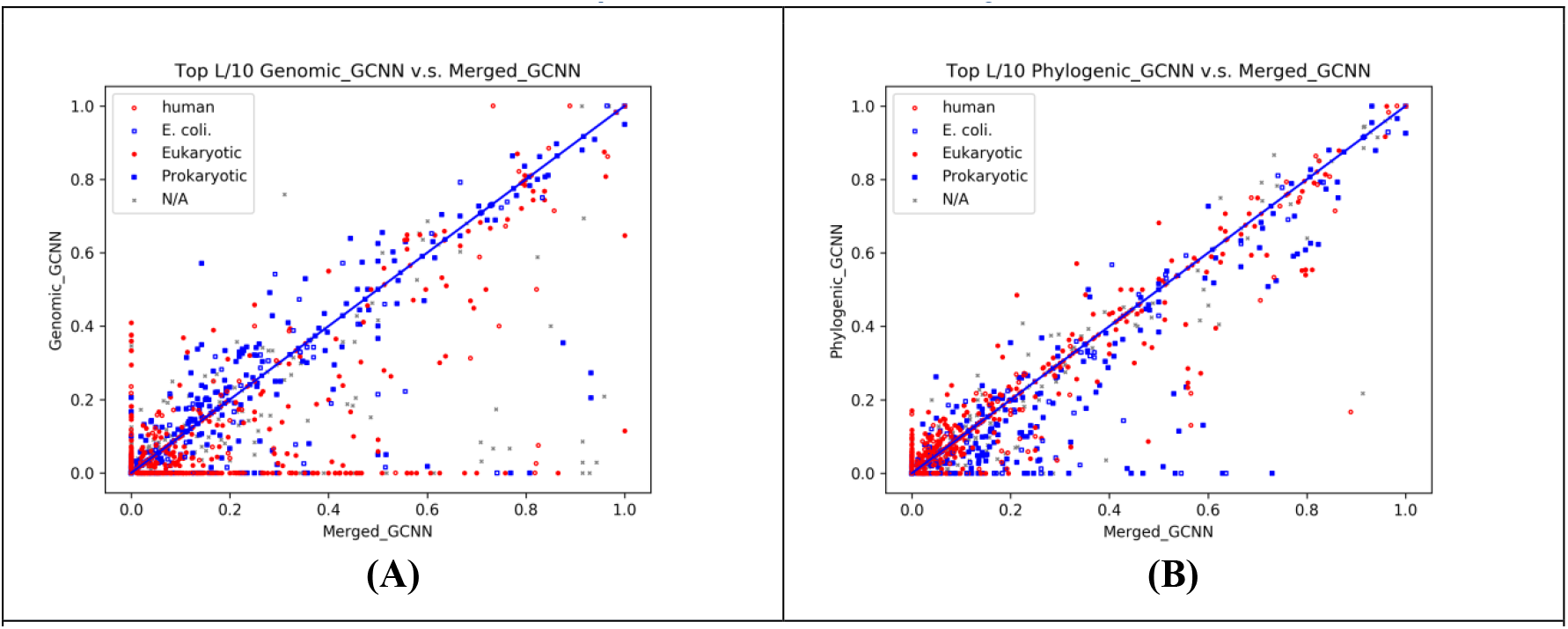
The top L/10 prediction accuracy of our DL method on MSAs generated by different methods. (A) merged (X-axis) vs. genome-based method (Y-axis); (B) merged (X-axis) vs. phylogeny-based method (Y-axis). Each dot represents one test protein pair in 3DComplex. All the 3DComplex protein pairs are colored by their species. Meanwhile, N/A indicates species other than eukaryotes and prokaryotes.

## 3. Specific example: study of an E. coli QFR complex

The quinol-fumarate reductase (QFR) respiratory complex (PDB ID 1KF6) of Escherichia coli is a four-subunit integral-membrane complex that catalyzes the final step of anaerobic respiration when fumarate is the terminal electron acceptor. The two chains, kf6_C and 1kf6_D, have 130 and 119 residues, respectively, and in total there are 146 contacts across them. Their individual MSAs have Meff ~72 and ~430, respectively. The paired MSAs created by the phylogeny- and genome-based methods have Meff ~60 and ~63, respectively. For a protein pair with such a small number of interlogs, our DL approach yields significantly better contact prediction than co-evolution methods including CCMpred, EVfold and Gremlin. As shown in Table S1, our method has top L/10 accuracy 79.17%, while CCMpred-, EVfold-, and Gremlin have top L/10 accuracy only 16.67%, 12.50%, and 20.83%, respectively. Figure S3 shows top L/10 predicted contact by each method and their overlaps with all the native contacts. This figure shows that CCMpred, EVfold and Gremlin predict some wrong contacts which are far away from any natives.

**Table S1.** Interfacial contact prediction accuracy (%) of our method GCNN, CCMpred, EVfold and Gremlin on the quinol-fumarate reductase (QFR) respiratory complex of E. coli (PDB: 1kf6).

| Prediction | L/5 | L/10 | L/20 | 50 | 20 | 10 | 5 |
| --- | --- | --- | --- | --- | --- | --- | --- |
| **GCNN** | 63.27 | 79.17 | 83.33 | 64.00 | 85.00 | 80.00 | 95.00 |
| **CCMpred** | 12.24 | 16.67 | 16.67 | 12.00 | 15.00 | 10.00 | 13.00 |
| **EVfold** | 6.12 | 12.50 | 16.67 | 6.00 | 15.00 | 20.00 | 17.00 |
| **Gremlin** | 14.29 | 20.83 | 16.67 | 14.00 | 20.00 | 20.00 | 21.00 |

**Figure S3.**
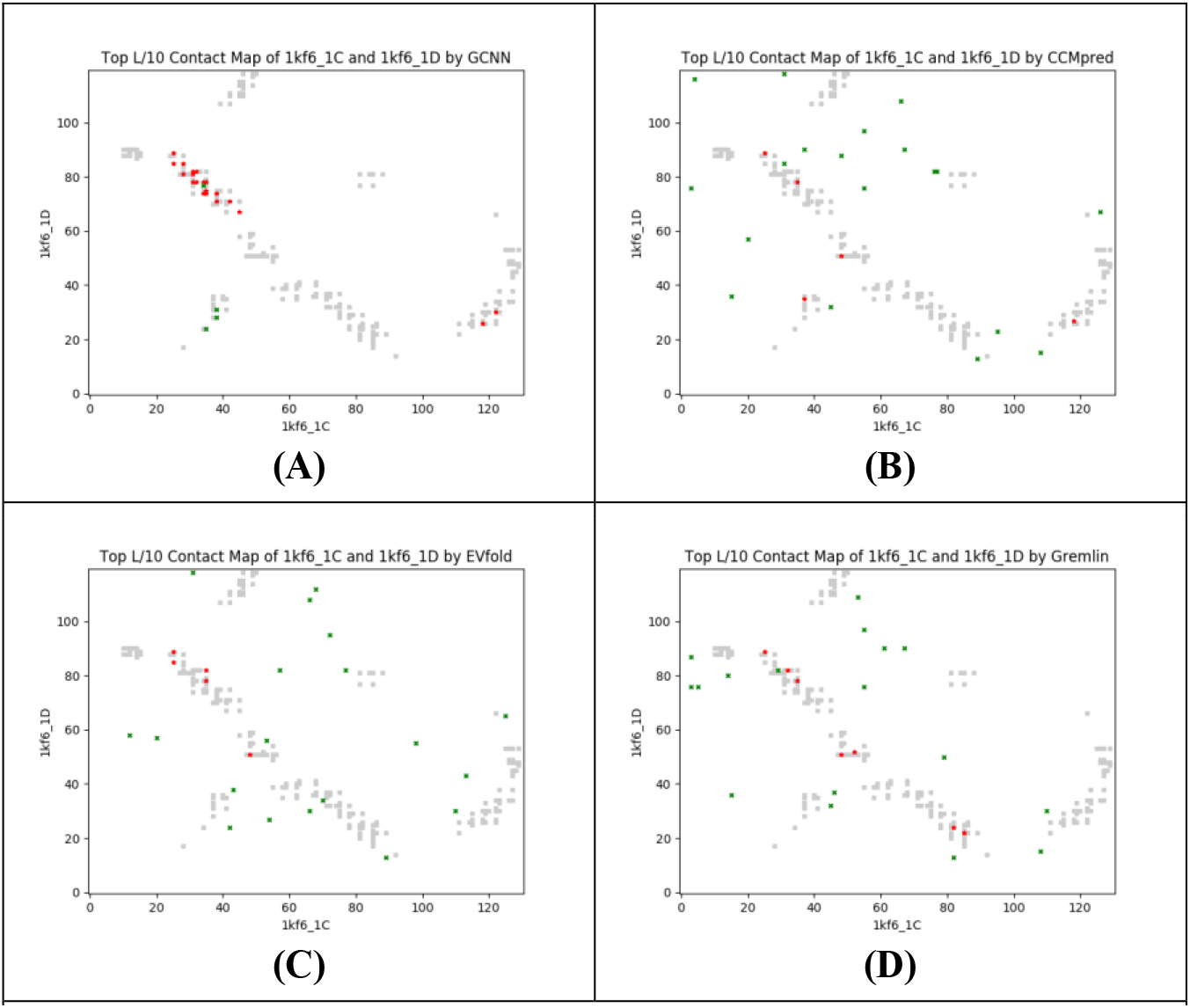
Overlap between the top L/10 predicted contacts (in red and green) and all the natives (in grey). Red (green) dots indicate correct (incorrect) predictions by (**A**) our approach, (**B**) CCMpred, (**C**) EVfold, and (**D**) Gremlin.

## References

1. Jones, S. and J.M. Thornton, Principles of protein-protein interactions. Proceedings of the National Academy of Sciences, 1996. 93(1): p. 13–20.

2. Sharan, R., et al., Conserved patterns of protein interaction in multiple species. Proceedings of the National Academy of Sciences of the United States of America, 2005. 102(6): p. 1974–1979.

3. Milenković, T., et al., Dominating biological networks. PloS one, 2011. 6(8): p. e23016.

4. Davis, D., et al., Topology-function conservation in protein–protein interaction networks. Bioinformatics, 2015. 31(10): p. 1632–1639.

5. Tuller, T., et al., Common and specific signatures of gene expression and protein–protein interactions in autoimmune diseases. Genes and immunity, 2013. 14(2): p. 67–82.

6. Lan, A., M. Ziv-Ukelson, and E. Yeger-Lotem, A context-sensitive framework for the analysis of human signalling pathways in molecular interaction networks. Bioinformatics, 2013. 29(13): p. i210–i216.

7. Yen, E.A., et al., Exploration of the dynamic properties of protein complexes predicted from spatially constrained protein-protein interaction networks. PLoS computational biology, 2014. 10(5): p. e1003654.

8. Deng, M., et al., Inferring domain–domain interactions from protein–protein interactions. Genome research, 2002. 12(10): p. 1540–1548.

9. Pritykin, Y. and M. Singh, Simple topological features reflect dynamics and modularity in protein interaction networks. PLoS computational biology, 2013. 9(10): p. e1003243.

10. Colak, R., et al. Dense graphlet statistics of protein interaction and random networks. in Pacific Symposium on Biocomputing. 2009.

11. Hormozdiari, F., et al., Protein-protein interaction network evaluation for identifying potential drug targets. Journal of Computational Biology, 2010. 17(5): p. 669–684.

12. Pržulj, N. and N. Malod-Dognin, Network analytics in the age of big data. Science, 2016. 353(6295): p. 123–124.

13. Lopes, A., et al., Protein-protein interactions in a crowded environment: an analysis via cross-docking simulations and evolutionary information. PLoS computational biology, 2013. 9(12): p. e1003369.

14. Dittrich, M.T., et al., Identifying functional modules in protein–protein interaction networks: an integrated exact approach. Bioinformatics, 2008. 24(13): p. i223–i231.

15. Cao, M., et al., Going the distance for protein function prediction: a new distance metric for protein interaction networks. PloS one, 2013. 8(10): p. e76339.

16. Ovchinnikov, S., H. Kamisetty, and D. Baker, Robust and accurate prediction of residue–residue interactions across protein interfaces using evolutionary information. Elife, 2014. 3: p. e02030.

17. Lensink, M.F., S. Velankar, and S.J. Wodak, Modeling protein-protein and protein-peptide complexes: CAPRI 6th edition. Proteins, 2017. 85(3): p. 359–377.

18. Mosca, R., A. Céol, and P. Aloy, Interactome3D: adding structural details to protein networks. Nature methods, 2013. 10(1): p. 47–53.

19. Marks, D.S., T.A. Hopf, and C. Sander, Protein structure prediction from sequence variation. Nat Biotechnol, 2012. 30(11): p. 1072–80.

20. Wang, S., et al., Accurate de novo prediction of protein contact map by ultra-deep learning model. PLoS computational biology, 2017. 13(1): p. e1005324.

21. Wang, S., et al., Folding membrane proteins by deep transfer learning. Cell Systems, 2017. 5(3): p. 202–211. e3.

22. Wang, S., S. Sun, and J. Xu, Analysis of deep learning methods for blind protein contact prediction in CASP12. Proteins: Structure, Function, and Bioinformatics, 2017.

23. Kosciolek, T. and D.T. Jones, De novo structure prediction of globular proteins aided by sequence variation-derived contacts. PLoS One, 2014. 9(3): p. e92197.

24. Wang, S., et al., CoinFold: a web server for protein contact prediction and contact-assisted protein folding. Nucleic acids research, 2016. 44(W1): p. W361–W366.

25. Hopf, T.A., et al., Sequence co-evolution gives 3D contacts and structures of protein complexes. Elife, 2014. 3: p. e03430.

26. Yu, J., et al., Lessons from (co-)evolution in the docking of proteins and peptides for CAPRI Rounds 28-35. Proteins, 2017. 85(3): p. 378–390.

27. Pellegrini, M., et al., Assigning protein functions by comparative genome analysis: protein phylogenetic profiles. Proceedings of the National Academy of Sciences, 1999. 96(8): p. 4285–4288.

28. Kortemme, T. and D. Baker, Computational design of protein–protein interactions. Current opinion in chemical biology, 2004. 8(1): p. 91–97.

29. Dandekar, T., et al., Conservation of gene order: a fingerprint of proteins that physically interact. Trends in biochemical sciences, 1998. 23(9): p. 324–328.

30. Galperin, M.Y. and E.V. Koonin, Who’s your neighbor? New computational approaches for functional genomics. Nature biotechnology, 2000. 18(6): p. 609–613.

31. Singh, R., et al., Struct2Net: a web service to predict protein–protein interactions using a structure-based approach. Nucleic acids research, 2010. 38(suppl_2): p. W508–W515.

32. Jothi, R., et al., Co-evolutionary analysis of domains in interacting proteins reveals insights into domain–domain interactions mediating protein–protein interactions. Journal of molecular biology, 2006. 362(4): p. 861–875.

33. Kim, E.D., et al., Predicting direct protein interactions from affinity purification mass spectrometry data. Algorithms for Molecular Biology, 2010. 5(1): p. 34.

34. Seemayer, S., M. Gruber, and J. Söding, CCMpred—fast and precise prediction of protein residue–residue contacts from correlated mutations. Bioinformatics, 2014. 30(21): p. 3128–3130.

35. Jones, D.T., et al., PSICOV: precise structural contact prediction using sparse inverse covariance estimation on large multiple sequence alignments. Bioinformatics, 2012. 28(2): p. 184–90.

36. de Juan, D., F. Pazos, and A. Valencia, Emerging methods in protein co-evolution. Nat Rev Genet, 2013. 14(4): p. 249–61.

37. Weigt, M., et al., Identification of direct residue contacts in protein-protein interaction by message passing. Proc Natl Acad Sci U S A, 2009. 106(1): p. 67–72.

38. Ekeberg, M., T. Hartonen, and E. Aurell, Fast pseudolikelihood maximization for direct-coupling analysis of protein structure from many homologous amino-acid sequences. Journal of Computational Physics, 2014. 276: p. 341–356.

39. Ma, J., et al., Protein contact prediction by integrating joint evolutionary coupling analysis and supervised learning. Bioinformatics, 2015. 31(21): p. 3506–3513.

40. Gueudré, T., et al., Simultaneous identification of specifically interacting paralogs and interprotein contacts by direct coupling analysis. Proceedings of the National Academy of Sciences, 2016. 113(43): p. 12186–12191.

41. Iserte, J., et al., I-COMS: Interprotein-COrrelated Mutations Server. Nucleic Acids Res, 2015. 43(W1): p. W320–5.

42. Rodriguez-Rivas, J., et al., Conservation of coevolving protein interfaces bridges prokaryote–eukaryote homologies in the twilight zone. Proceedings of the National Academy of Sciences, 2016. 113(52): p. 15018–15023.

43. Levy, E.D., et al., 3D complex: a structural classification of protein complexes. PLoS computational biology, 2006. 2(11): p. e155.

44. Remmert, M., et al., HHblits: lightning-fast iterative protein sequence searching by HMM-HMM alignment. Nature methods, 2012. 9(2): p. 173–175.

45. Feinauer, C., et al., Inter-protein sequence co-evolution predicts known physical interactions in bacterial ribosomes and the Trp operon. PloS one, 2016. 11(2): p. e0149166.

46. Zhu, H., M. Zhou, and R. Alkins, Group role assignment via a Kuhn–Munkres algorithm-based solution. IEEE Transactions on Systems, Man, and Cybernetics-Part A: Systems and Humans, 2012. 42(3): p. 739–750.

47. Pakseresht, N., et al., Assembly information services in the European Nucleotide Archive. Nucleic acids research, 2013. 42(D1): p. D38–D43.

48. Federhen, S., The NCBI taxonomy database. Nucleic acids research, 2011. 40(D1): p. D136–D143.

49. Wang, S., et al., Protein secondary structure prediction using deep convolutional neural fields. Scientific reports, 2016. 6.

50. Ma, J. and S. Wang, AcconPred: Predicting solvent accessibility and contact number simultaneously by a multitask learning framework under the conditional neural fields model. BioMed research international, 2015. 2015.

51. Wang, S., et al., RaptorX-Property: a web server for protein structure property prediction. Nucleic acids research, 2016. 44(W1): p. W430–W435.

52. He, K., et al., Deep residual learning for image recognition. arXiv preprint arXiv:1512.03385, 2015.

53. De Juan, D., F. Pazos, and A. Valencia, Emerging methods in protein co-evolution. Nature Reviews Genetics, 2013. 14(4): p. 249–261.

